# The molecular mechanism of uptake and cell-to-cell transmission of arginine-containing dipeptide repeat proteins

**DOI:** 10.1101/2025.11.25.690484

**Authors:** Alexandra B. Sutter, Benito F. Buksh, Jelena Mojsilovic-Petrovic, Casey Dalton, Nicholas A. Till, Danielle C. Morgan, David W. C. MacMillan, Robert G. Kalb

## Abstract

Micro-satellite repeat expansion of the 5’ GGGGCC 3’ sequence in the *C9orf72* gene is the most common monogenic form of amyotrophic lateral sclerosis (ALS) and frontotemporal dementia (FTD). Dipeptide repeat proteins (DPRs) translated from the mutant allele can be detected in postmortem brains of afflicted individuals. The arginine containing peptides, poly-PR and poly-GR, are particularly noxious to cells. Both have been shown to undergo cell-cell transmission, but the underlying mechanisms are not understood. We found rapid internalization and nucleolar localization of bath-applied hemagglutinin (HA) tagged poly-PR with twenty repeats (HA-PR_20_) in cell lines and neurons. Small molecule and RNAi approaches implicated a temperature-dependent, fluid phase endocytosis mechanism in HA-PR_20_ uptake. We sought to identify DPR-related cell surface uptake factors using a high-resolution proximity labeling technique developed in the MacMillan group, termed µMap. DPR-iridium conjugates identified candidate cell-surface proteins which were interrogated in an RNAi screen. Focusing on our strongest candidate, chondroitin sulfate proteoglycan 4 (CSPG4), we showed that cellular uptake of HA-PR_20_ is blocked by inhibition of glycosaminoglycan chain synthesis (using drugs or RNAi) and knockdown or ablation of CSPG4 (using RNAi or CRISPR editing). Reduction of CSPG4 protected PR_20_-induced neuronal toxicity. We used a dual reporter system to interrogate *in vitro* neuron-to-neuron transmission of PR_50_ and found that PR_50_ synthesized by one neuron readily spread to neighboring neurons. Transmission was significantly reduced when CSPG4 was knocked down. These results suggest CSPG4 is an important factor in poly-PR internalization and transmission and therefore may be a therapeutic target to slow DPR transmission and disease progression.

**Significance Statement:** A GGGGCC hexanucleotide repeat expansion in the *C9orf72* gene is the most common monogenic form of ALS/FTD. This expansion leads to dipeptide repeat protein (DPR) production through non-canonical translation of repeat-containing RNA. DPRs have been shown to transmit between cells, but how this occurs is not well understood. We identified the cell surface protein chondroitin sulfate proteoglycan 4 (CSPG4) as a mediator of the uptake and intercellular spread of toxic arginine-rich DPRs. Targeting CSPG4 may provide a strategy to block DPR transmission and slow disease progression.

## Introduction

An intronic GGGGCC hexanucleotide repeat expansion in the *chromosome 9 open reading frame 72 (C9orf72)* gene is the most common monogenic form of amyotrophic lateral sclerosis (ALS) and frontotemporal dementia (FTD), accounting for approximately 25-40% of familial cases (1–3). Affected individuals typically harbor hundreds to thousands of these GGGGCC repeats (4, 5). Three primary pathogenic mechanisms have been proposed: (1) loss of C9orf72 function due to haploinsufficiency (6–8); (2) RNA toxicity from expanded repeat-containing transcripts sequestering RNA binding proteins (1, 9–19); and (3) toxicity from dipeptide repeat proteins (DPRs) generated via repeat-associated non-ATG translation (RAN-T) (17, 19–24). The precise contribution of each mechanism to disease pathogenesis remains unresolved.

DPR pathology emerges prior to other key pathological features of ALS, including TDP-43 aggregation (25–28). A neuropathological hallmark of *C9orf72*-mediated disease is the presence of cytoplasmic inclusions in both neurons and glial cells that are negative for TDP-43, but positive for p62, ubiquitin, and the five DPR species produced through RAN-T (20, 29, 30). Among these, the arginine-rich DPRs – poly-PR and poly-GR – are highly polar and exhibit pronounced toxicity *in vitro* and *in vivo* (24, 31–37).

In post-mortem brain tissue from patients with *C9orf72*-associated ALS, DPRs are found in aggregates distributed across multiple regions (29, 38, 39). These aggregates appear both in isolated cells and in clusters, suggesting focal sites of active RAN-T and possibly reflecting local propagation of pathology (40). One proposed mechanism of disease progression in *C9orf72*-associated ALS, as in other neurodegenerative diseases, is the transcellular spread of pathogenic proteins (41–43). Misfolded and aggregated proteins may transit between cells, seeding the nucleation and aggregation of soluble counterparts in recipient cells, as has been described for TDP-43 (44–52), tau (53, 54), and α-synuclein (47, 55, 56). This process can explain the spread of pathology across anatomically connected brain regions, thereby contributing to disease progression. All five DPR species have been shown to undergo cell-to-cell transmission *in vitro* (42), although the molecular mechanisms mediating their intercellular transfer remain poorly understood.

In this study, we sought to elucidate the mechanisms underlying intercellular transmission of DPRs, with a particular focus on the arginine-rich species. We found that poly-PR is internalized by recipient cells via an energy-dependent mechanism involving macropinocytosis. To identify the molecular basis for poly-PR uptake, we used a proximity labeling approach. This powerful strategy for interrogating protein interactions within native cellular contexts has recently been leveraged to map trafficking pathways of cell-penetrating peptides (CPPs) in live cells (57, 58). Using µMap proximity labeling, we identified chondroitin sulfate proteoglycan 4 (CSPG4) as a prominent cell-surface protein implicated in DPR uptake. We further demonstrated that CSPG4 is a critical mediator of arginine-rich DPR internalization and intercellular spread. CSPG4 knockdown significantly reduced both poly-PR uptake and its transmission between cells and conferred neuroprotection. Together, these findings implicate CSPG4 as a potential therapeutic target to limit DPR propagation and slow disease progression in C9orf72-associated ALS, and FTD or associated diseases.

## Results

### PR_20_ uptake pathway shares features with macropinocytosis

To investigate how arginine-rich DPRs are internalized by cells, we started with a simplified assay monitoring uptake of bath-applied hemagglutinin (HA)-tagged poly-PR consisting of 20 repeats (HA-PR_20_) to both HeLa cells and primary rat neuron cultures. Because PR is enriched at the cell surface after bath application, by western blot we did not detect an increase in uptake over time (Fig. 1A, Membrane +). To assess internalized peptide specifically, cell surface-bound PR_20_ was removed via trypsin digestion prior to analysis. This allowed us to monitor time-dependent uptake of HA-PR_20_, with robust internalization visible within minutes (Fig. 1A, Membrane –) and nucleolar localization apparent by 30 minutes post-application (Fig. 1B), both by western blot and immunocytochemistry (ICC). These results aligned with those from the McKnight lab (36, 59)

**Fig. 1.**
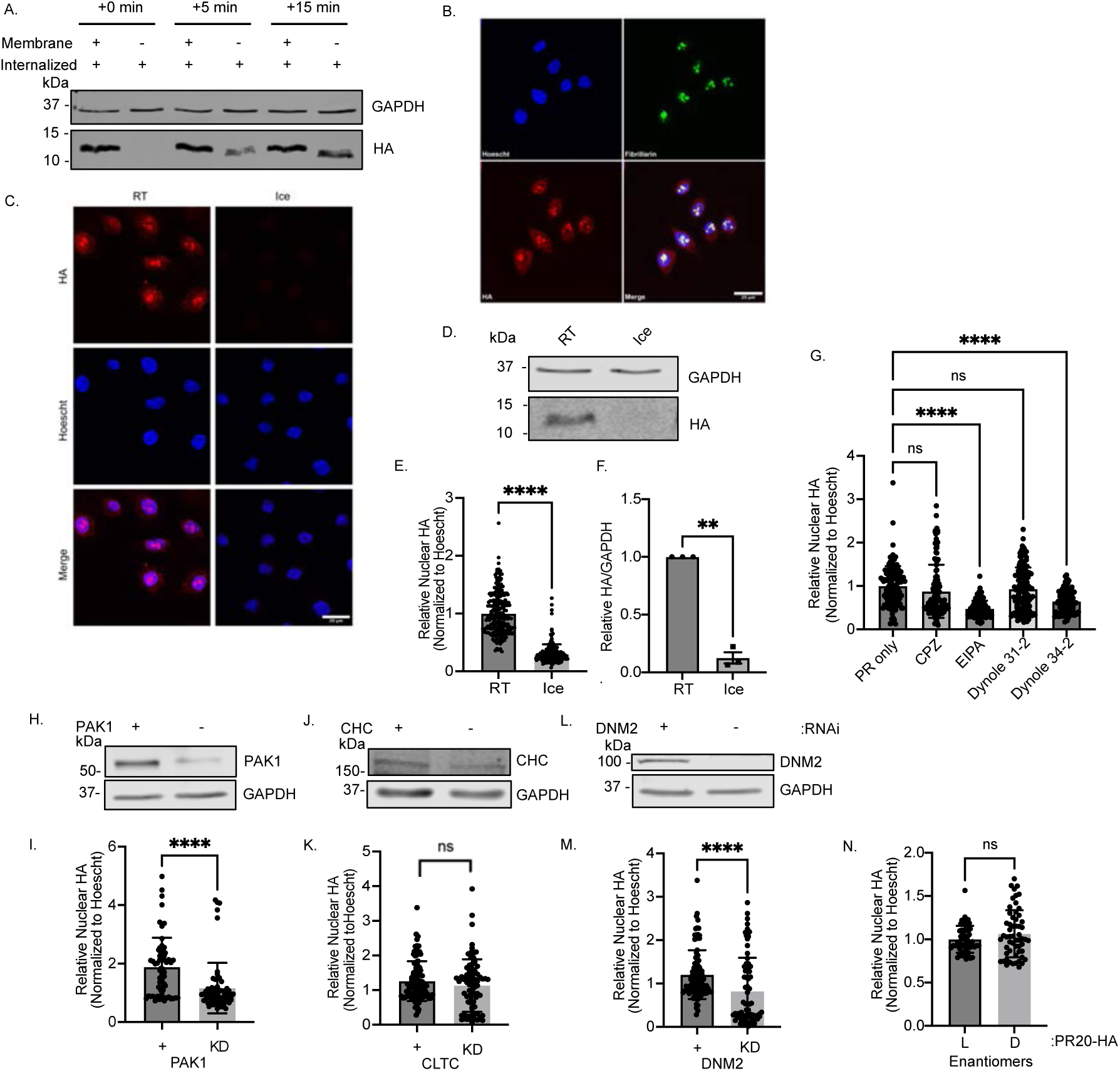
Uptake of PR_20_ is mediated by a pathway that shares features with macropinocytosis. (A) Western blot detection of HA-PR_20_ uptake over 15 minutes in HeLa cells. Membrane (+) fractions are membrane bound + internalized PR; (−) fractions are solely internalized peptide. GAPDH, loading control. (B) Representative ICC of HA-PR_20_ colocalization with fibrillarin, a nucleolar marker, after 30-minutes of bath application. Scale bar, 20µm. Blue = Hoescht, green = fibrillarin, red = HA. (C) Representative ICC of HA-PR_20_ bath application at room temperature or on ice. Scale bar, 20µm. Blue = Hoescht, Red = HA. (D) Representative western blot of internalized HA-PR_20_ after bath application at room temperature or on ice. GAPDH, loading control. (E) Quantification of (C). Unpaired *t* test, two-tailed, n=3 biological replicates, ****p<0.0001. (F) Quantification of (E). Unpaired *t* test, two-tailed, n=3 biological replicates, **p<0.01 (G) Quantification of internalized HA-PR_20_ in neurons pretreated with endocytosis inhibitors (chlorpromazine (CPZ), clathrin-mediated endocytosis inhibitor; 5-(N-ethyl-N-isopropyl)amiloride (EIPA), macropinocytosis inhibitor; and dynamin inhibitor Dynole 34-2 (inactive isomer Dynole 31-2)) for 1 hour before a 1-hour bath application with HA-PR_20_. One way ANOVA with Tukey’s *post hoc* test, F (_5,728_) = 18.02, n=3 biological replicates, ****p<0.0001. (H) Representative western blot showing p21-activated kinase 1 (PAK1) knockdown in HeLa cells. GAPDH, loading control. (I) Quantification of nuclear HA-PR_20_ after PAK1 KD in HeLa cells. Unpaired *t* test, two-tailed, n=3 biological replicates, ****p<0.0001. (J) Representative western blot of clathrin heavy chain (CHC) knockdown in HeLa cells. GAPDH, loading control. (K) Quantification of nuclear HA-PR_20_ after CHC KD in HeLa cells. Unpaired *t* test, two-tailed, n=3 biological replicates, ns. (L) Representative western blot of dynamin 2 (DNM2) knockdown in HeLa cells. GAPDH, loading control. (M) Quantification of nuclear HA-PR_20_ after DNM2 KD in HeLa cells. Unpaired *t* test, two-tailed, n=3 biological replicates, ****p<0.0001. (N) Quantification of ICC of 30-minute bath application of HeLa with HA-PR_20_ composed of either D-amino acids or L-amino acids. Unpaired *t* test, two-tailed, n=3 biological replicates, ns.

We next sought to determine whether poly-PR uptake occurs via endocytosis or direct translocation across the plasma membrane. Since endocytosis is an energy-dependent process that is inhibited at low temperatures, we applied HA-PR_20_ to cells at either room temperature or at 4 °C for 5 minutes, followed by a wash and then assay. PR_20_ was readily internalized at room temperature by HeLa cells, but markedly reduced at 4 °C, as shown by both immunocytochemistry and western blot analysis (Fig. 1C–F). These results suggested that poly-PR internalization is energy-dependent.

To determine the specific endocytic pathway involved, we pretreated HeLa cells with various endocytosis inhibitors one hour prior to HA-PR_20_ application and assessed uptake by immunocytochemistry and western blotting. Nuclear accumulation of HA-PR_20_ was significantly reduced in cells pretreated with 5-(*N*-ethyl-*N*-isopropyl)amiloride (EIPA), an inhibitor of macropinocytosis (Fig. S1A), with chlorpromazine (CPZ), an inhibitor of clathrin-mediated endocytosis (Fig. S1B), and the dynamin inhibitor Dynole 34-2 (but not its inactive isomer Dynole 31-2) (Fig. S1C). These results were also seen in primary mixed spinal cord cultures, however CPZ pretreatment did not significantly reduce PR uptake (Fig. 1G). Western blot analysis following pulse-chase experiments revealed a significant reduction in PR_20_ uptake only in the presence of EIPA (Fig. S1D-G), suggesting that macropinocytosis is the predominant route of internalization.

To confirm the specificity of these results and account for potential off-target effects of pharmacological inhibitors, we used siRNA-mediated knockdown of key endocytic regulators: clathrin heavy chain (CLTC), dynamin 2 (DNM2), and p21-activated kinase 1 (PAK1) (60). Knockdown of DNM2 and PAK1, but not CLTC, significantly reduced nuclear HA-PR_20_ signal relative to scrambled siRNA controls (Fig. 1H-M), further supporting a role for dynamin-dependent macropinocytosis in PR_20_ uptake.

Finally, to assess whether the uptake mechanism depends on peptide stereochemistry, we synthesized and applied HA-PR_20_ composed of D-amino acids. Internalization of D-PR_20_ did not significantly differ from that of L-PR_20_ (Fig. 1N), consistent with a non-receptor-mediated, fluid-phase uptake mechanism.

Taken together, these data indicated that arginine-rich DPRs such as poly-PR are internalized through a dynamin-dependent pathway primarily mediated by macropinocytosis.

### µMap reveals candidate cell surface molecules mediating uptake of PR_20_

To identify the specific cell surface proteins that facilitate PR_20_ internalization, we employed µMap, an iridium-based photocatalytic proximity labeling platform. This method leverages Dexter energy transfer to generate reactive carbenes from co-applied biotin-diazirines, enabling covalent biotinylation of proteins within a ∼1 nm radius of the iridium-tagged molecule. Proximal biotinylated proteins are identified downstream by proteomics analysis (61).

We applied µMap to label proteins in proximity to poly-PR at the cell-surface. Iridium-tagged PR_20_ (Ir-PR_20_) (Fig. S2B-C) was applied to HeLa cells on ice to prevent internalization and restrict labeling to the membrane. After 5 minutes of incubation on ice, biotin-diazirine (Fig. S2A) was added and blue light was applied to label proteins in proximity to Ir-PR_20_ at the cell surface. In parallel, we compared this to cells similarly pulsed with Ir-PR_20_ followed by a 1 hour chase at 37 °C to allow internalization. HA-PR_20_ signal was enriched on the membrane only during the surface-bound pulse phase, as demonstrated by co-localization with Wheat Germ Agglutinin (WGA) (Fig. 2A-B).

**Fig. 2.**
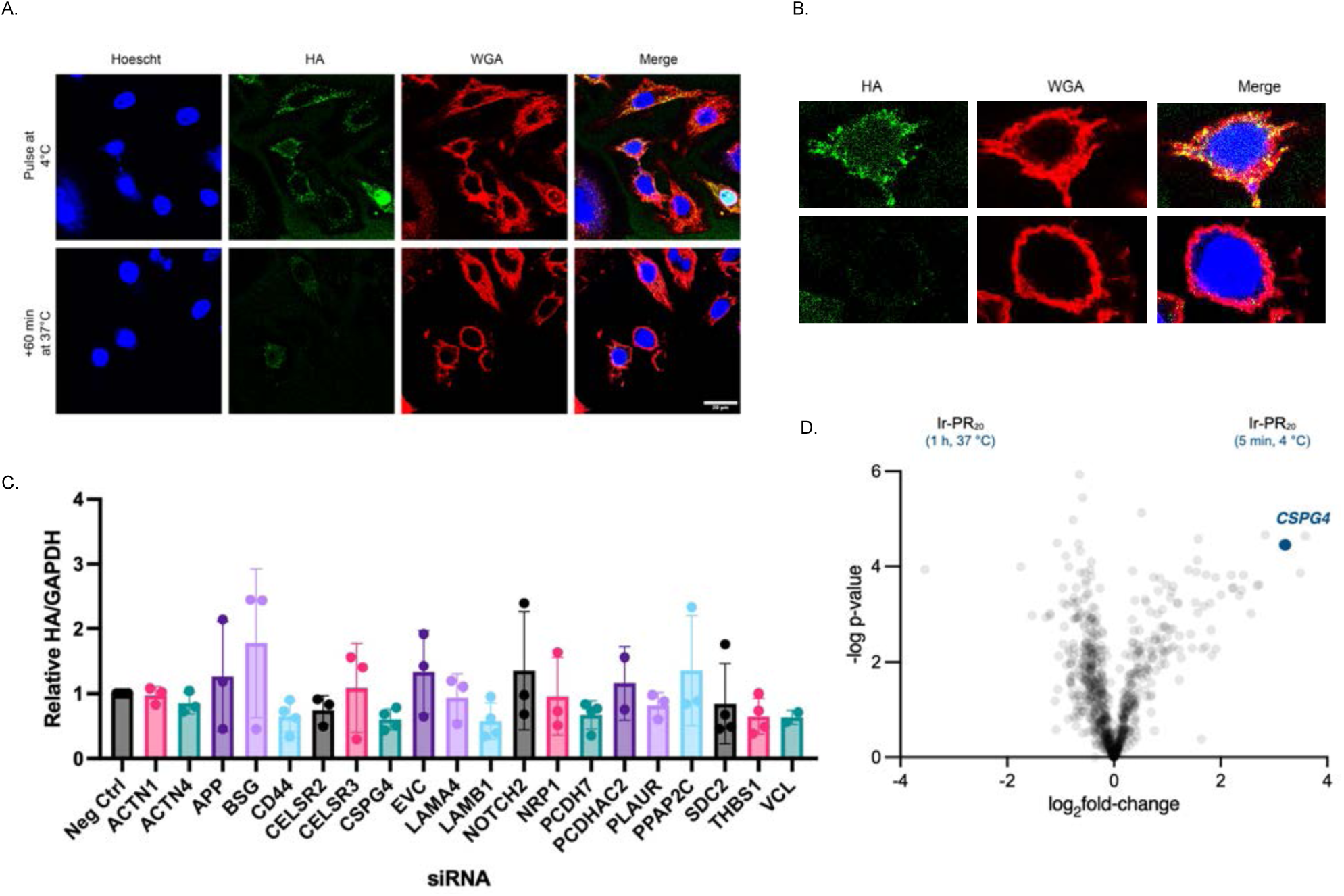
µMap proximity labeling followed by mass spectrometry reveals cell surface candidates for PR_20_ uptake. (A). Representative ICC of µMap conditions. After a 5-minute pulse on ice, HA-PR_20_ is localized to the cell surface, as labeled with Wheat Germ Agglutinin (WGA). With a subsequent 60-minute chase at 37°C, HA-PR_20_ is no longer enriched on the cell surface. Single z-stack, scale bar, 20µm. Blue = Hoescht, Green = HA, Red = WGA. (B) Enlarged view of cells in (A). (C) Quantification of siRNA pulse/chase screen. HeLa cells with siRNA knockdown of indicated genes were treated with a 5-minute HA-PR_20_ pulse followed by a 5-minute chase. Internalized peptide was obtained and immunoblots were performed for HA. GAPDH, loading control. Quantification by normalizing HA signal to GAPDH signal. n=3 biological replicates. (D) Volcano plot of µMap mass spectrometry results. CSPG4 is highlighted.

Proteomics analysis of labeled proteins yielded a list of candidates, which were filtered to twenty proteins based on two criteria: (1) expression in neurons, and (2) localization to the cell surface (Fig. S2D-E). First, we assessed their functional relevance by conducting an siRNA-based screen in HeLa cells using our established pulse-chase assay to quantify internalized HA-PR_20_ by western blotting. Knockdown of several candidates produced significant changes in HA-PR_20_ uptake compared to controls, with some proteins showing marked reductions in internalization (Fig. 2C).

Among the candidates, chondroitin sulfate proteoglycan 4 (CSPG4) emerged as the top hit, showing the most robust and reproducible reduction in poly-PR internalization. Consistent with this, CSPG4 appeared as one of the most enriched proteins in the µMap proteomics volcano plot (Fig. 2D). Given that under physiological conditions, chondroitin sulfate proteoglycans are negatively charged and arginine-rich DPRs are positively charged, the electrostatic interaction between CSPG4 and poly-PR provided a compelling mechanistic rationale for its involvement in mediating uptake. These findings prompted us to further investigate CSPG4 as a potential uptake factor of poly-PR internalization.

### CSPG4 is an uptake factor for arginine-rich DPRs

To confirm that CSPG4 directly mediates poly-PR internalization, we first tested for physical interaction between HA-PR_20_ and CSPG4. Following a 30-minute bath application of HA-PR_20_, we detected interaction using proximity ligation assay in HEK (Fig. S3A).

We next investigated the role of CSPG4 as a PR uptake mediator either by targeting both its expression and its associated glycosaminoglycan (GAG) side chains. We validated CSPG4 knockdown in HeLa cells via siRNA transfection, confirming reduced protein levels by western blot relative to scrambled siRNA controls (Fig. 3A). In a pulse-chase assay, we observed an approximately 40% reduction in HA-PR_20_ uptake in CSPG4 KD cells via western blot (Fig. 3B–C). Additionally, enzymatic digestion of chondroitin sulfate proteoglycan side chains with Chondroitinase ABC or knockdown of xylosyltransferase 2 (XYLT2), an enzyme responsible for initiating GAG chain addition to proteoglycans, led to significant ∼50% decreases in poly-PR internalization compared to controls (Fig. S3B-E), indicating that chondroitin sulfate (CS) chains are likely to be critical for this process.

**Fig. 3.**
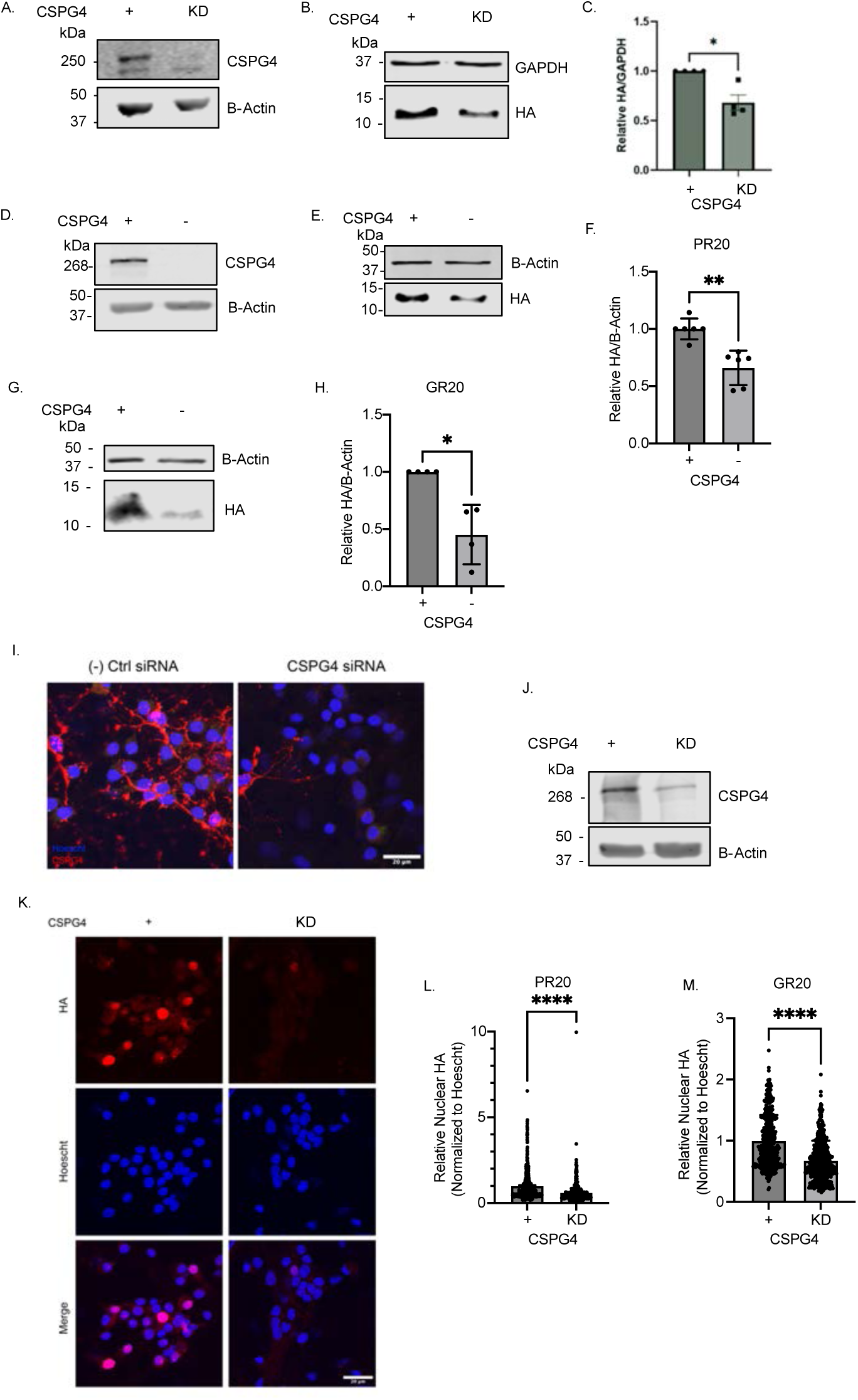
Arginine-rich DPR uptake is significantly reduced by reducing CSPG4 abundance. (A) Representative western blot of CSPG4 knockdown in HeLa cells. B-Actin, loading control. (B) Representative western blot of a 5-minute HA-PR_20_ pulse followed by a 5-minute chase. Internalized peptide was compared in negative control siRNA conditions relative to CSPG4 knockdown conditions. (C) Quantification of HA-PR_20_ internalization. HA was normalized to B-Actin loading control. Unpaired *t* test, two-tailed, n=4 biological replicates, *p<0.05. (D) Representative western blot of CSPG4 levels in Flp-In T-REx HEK CSPG4-knockout (CSPG4⁻^/^⁻) cells. B-Actin, loading control. (E) Representative western blot of a 5-minute HA-PR_20_ pulse followed by a 15-minute chase in CSPG4^+/+^ and CSPG4⁻^/^⁻ cells. B-Actin, loading control. (F) Quantification of (E). Unpaired *t* test, two-tailed, n=5 biological replicates, **p<0.01. (G) Representative western blot of a 5-minute HA-GR_20_ pulse followed by a 15-minute chase in CSPG4^+/+^ and CSPG4^−/−^ cells. B-Actin, loading control. (H) Quantification of (G). Unpaired *t* test, two-tailed, n=3 biological replicates, *p<0.05. (I) Representative western blot of CSPG4 siRNA knockdown in primary rat mixed spinal cord cultures. B-Actin, loading control. (J) Representative ICC of CSPG4 siRNA knockdown in primary rat mixed spinal cord cultures. Scale bar, 20 µm. Blue = Hoescht, Red = CSPG4. (K) Representative 2-hour HA-PR_20_ bath application in primary mixed spinal cord cultures with negative control and CSPG4 siRNA knockdown conditions. Scale bar, 20 µm. Blue = Hoescht, Red = HA. (L) Quantification of (K). Unpaired *t* test, two-tailed, n=3 biological replicates, ****p<0.001. (M) Quantification of 2-hour HA-GR_20_ bath application in primary mixed spinal cord cultures with negative control and CSPG4 siRNA knockdown conditions. Unpaired *t* test, two-tailed, n=3 biological replicates, ****p<0.001.

Next, we generated CSPG4-knockout (CSPG4⁻^/^⁻) cells using CRISPR-Cas9 in the Flp-In T-REx HEK cell system (Fig. 3D). In CSPG4⁻^/^⁻ cells, internalization of both HA-PR_20_ and HA-GR_20_ was significantly reduced compared to wild-type CSPG4⁺^/^⁺ cells in the pulse-chase paradigm (Fig. 3E-H). The Flp-In T-REx system enabled doxycycline-inducible CSPG4 overexpression in both CSPG4⁺^/^⁺ and CSPG4⁻^/^⁻ backgrounds. This allowed us to: (1) overexpress CSPG4 in wild-type cells, and (2) reintroduce CSPG4 into the knockout background (Fig. S3F-G). Reintroducing CSPG4 expression in KO cells restored the uptake of both HA-PR_20_ and HA-GR_20_, confirming CSPG4’s role in mediating internalization of arginine-rich DPRs (Fig. S3H-I, L-M). Overexpression of CSPG4 in wild-type cells did not further increase DPR uptake, suggesting that physiological levels of CSPG4 are limiting for uptake (Fig. S3J-K, N-O).

We next asked whether these findings translated to a neuronal context. To test this, we transfected primary mixed spinal cord cultures at 4 days *in vitro* (DIV4) with siRNA targeting CSPG4. By DIV14, we confirmed robust knockdown by both immunocytochemistry and western blot (Fig. 3I-J). HA-PR_20_ or HA-GR_20_ was bath-applied, and nuclear uptake was assessed via immunocytochemistry. In both conditions, we observed a significant reduction in nuclear HA signal relative to negative controls, indicating that CSPG4 is required for internalization of arginine-containing DPRs in neurons (Fig. 3K-M). Consistent with this, pretreatment of DIV14 neurons with Chondroitinase ABC similarly reduced nuclear uptake of both HA-PR_20_ and HA-GR_20_ (Fig. S3P-Q).

To pharmacologically inhibit CSPG biosynthesis, we treated DIV14 neurons with Fluorosamine (peracetylated 4-F-GlcNAc) or Difluorosamine (Ac-4,4-diF-GlcNAc), both inhibitors of GAG chain synthesis ((62, 63)), at 10 μM for 24 hours. Following peptide application, nuclear HA signal was significantly decreased in both treatment groups compared to vehicle controls, reinforcing the critical role of CSPGs and their GAG modifications in mediating neuronal uptake of arginine-rich DPRs (Fig. S3R-S).

### Blocking PR_20_ uptake is neuroprotective and influences TDP-43 proteinopathy

Having established CSPG4 as a key mediator of poly-PR internalization, we next investigated whether reducing CSPG4 expression could mitigate the toxic effects of arginine-rich DPRs in neurons. Given the known temporal relationship between DPR accumulation and the subsequent development of TDP-43 pathology, we asked whether exposure to poly-PR could induce formation of pathological, sarkosyl-insoluble TDP-43, a biochemical hallmark of many neurodegenerative diseases (64).

We bath-applied HA-PR_20_ to primary cortical neurons for 48 hours and detected the presence of sarkosyl-insoluble TDP-43 by western blot—an insoluble species absent in untreated controls—suggesting that poly-PR exposure alone is sufficient to trigger this pathological entity.

To assess whether CSPG4 modulates this effect, we transfected primary cortical neurons at DIV4 with siRNA targeting CSPG4, followed by a 48-hour treatment with HA-PR_20_ on DIV17. While levels of soluble TDP-43 remained unchanged, CSPG4 knockdown resulted in a significant reduction of sarkosyl-insoluble TDP-43 compared to scrambled siRNA controls (Fig. 4A–B). These findings suggested that CSPG4-mediated internalization of poly-PR contributes to the induction of insoluble TDP-43 species.

**Fig. 4.**
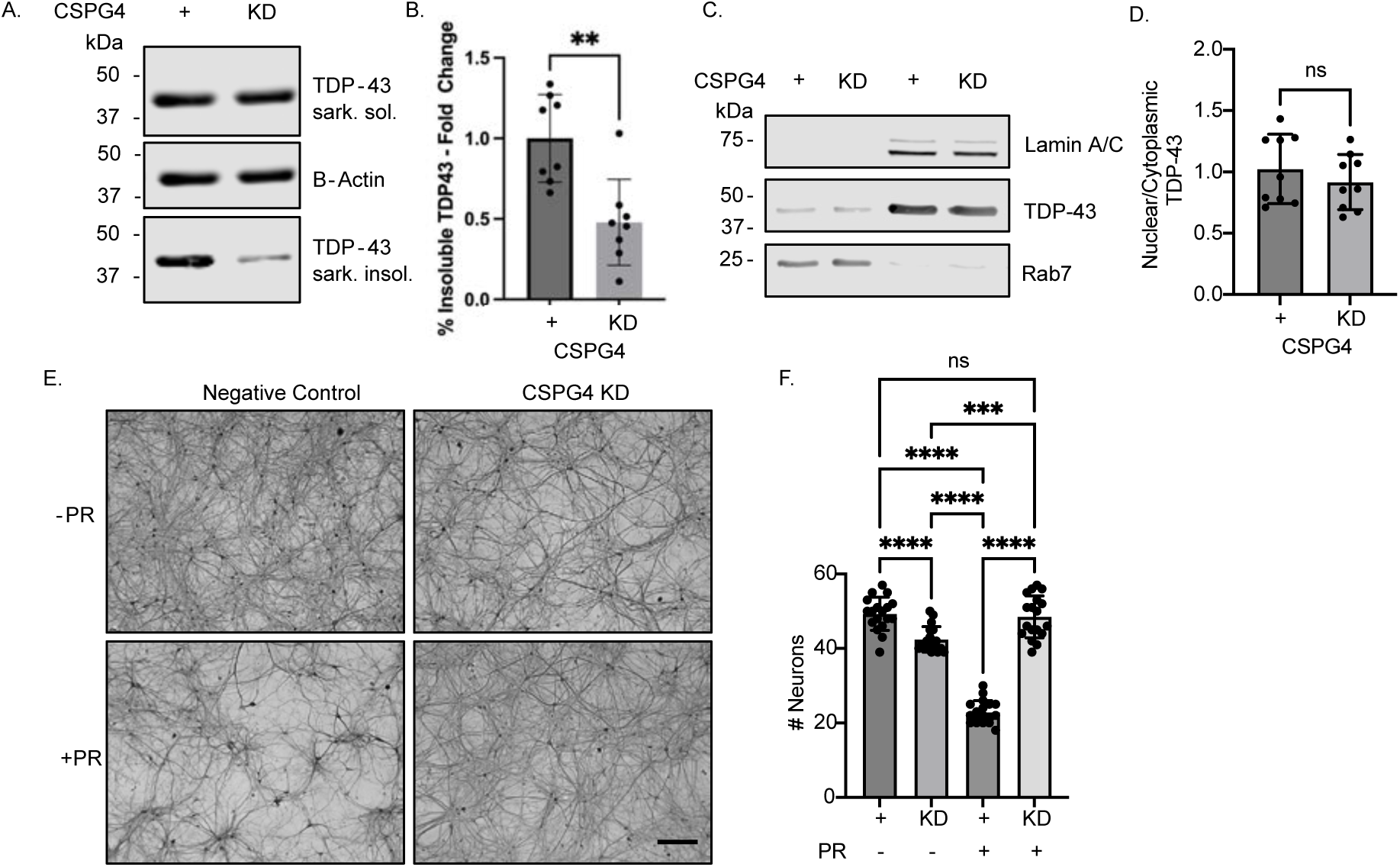
Reducing PR_20_ uptake reduces sarkosyl insoluble TDP-43 species and is neuroprotective. (A) Representative sarkospin western blot of negative control and CSPG4 siRNA knockdown rat primary cortical neurons treated with HA-PR_20_ for 48 hours. Immunoblotting for Sarkosyl soluble (sark. sol.) and insoluble (sark. insol.) TDP-43. B-Actin, loading control. (B) Quantification of (A). Unpaired *t* test, two-tailed, n=3 biological replicates, **p<0.01. (C) Representative western blot of nuclear and cytoplasmic fractionation in negative control and CSPG4 siRNA knockdown rat primary cortical neurons treated with HA-PR_20_ for 48 hours. Immunoblot for TDP-43. Lamin A/C, nuclear loading control. Rab7, cytoplasmic loading control. (D) Quantification of (C). Unpaired *t* test, two-tailed, n=4 biological replicates, ns. (E) Representative SMI-32 brightfield images of negative control and CSPG4 siRNA knockdown rat primary mixed spinal cord cultures treated with or without HA-PR_20_ for 5 days. Scale bar, 300 µm. (F) Quantification of (E). One way ANOVA with Tukey’s *post hoc* test, F (_3,68_) = 149.3, n=3 biological replicates, ****p<0.001.

We next asked whether TDP-43 mislocalization—another common feature of ALS pathology—was altered under these conditions. Surprisingly, we did not observe any changes in the nuclear-to-cytoplasmic distribution of TDP-43 by western blot of subcellular fractions at the 48-hour time point (Fig. 4C–D).

To examine whether CSPG4 knockdown confers neuroprotection in the presence of poly-PR, we turned to our primary mixed spinal cord culture model. At DIV14, we applied 2 μM HA-PR_20_ for 5 days. This treatment resulted in approximately 50% motor neuron loss, as assessed by SMI-32 staining, relative to untreated cultures as seen previously (65). Remarkably, CSPG4 knockdown fully rescued motor neuron survival, restoring levels comparable to untreated controls (Fig. 4E–F). These data indicated that CSPG4 is not only involved in DPR uptake but also contributes to downstream neurotoxicity.

### CSPG4 mediates intercellular spreading of PR pathology *in vitro*

To determine whether our findings extended to intercellular transmission of DPRs, we developed a model of neuron-to-neuron DPR spread. Unlike synthetic HA-PR_20_ peptide, which is exogenously applied and not synthesized by cells, DPRs in *C9orf72* ALS patients are generated intracellularly. To more accurately reflect this, we employed a dual-reporter adeno-associated virus (AAV) system to examine the cell-to-cell propagation of PR_50_ in primary neurons.

Neurons were transduced with an AAV encoding GFP-T2A-PR_50_-FLAG, which enables the co-expression of GFP and PR_50_-FLAG in the same cell. After 5 days, we confirmed robust expression of both GFP and PR_50_-FLAG in transduced neurons. Using immunocytochemistry, we observed PR_50_-FLAG puncta in neighboring, GFP-negative neurons, illustrating that DPRs synthesized by one neuron can spread to adjacent cells (Fig. 5A). These puncta were primarily cytoplasmic or perinuclear and were detected in both MAP2-positive and MAP2-negative cells, suggesting spread to both neurons and glia.

**Fig. 5.**
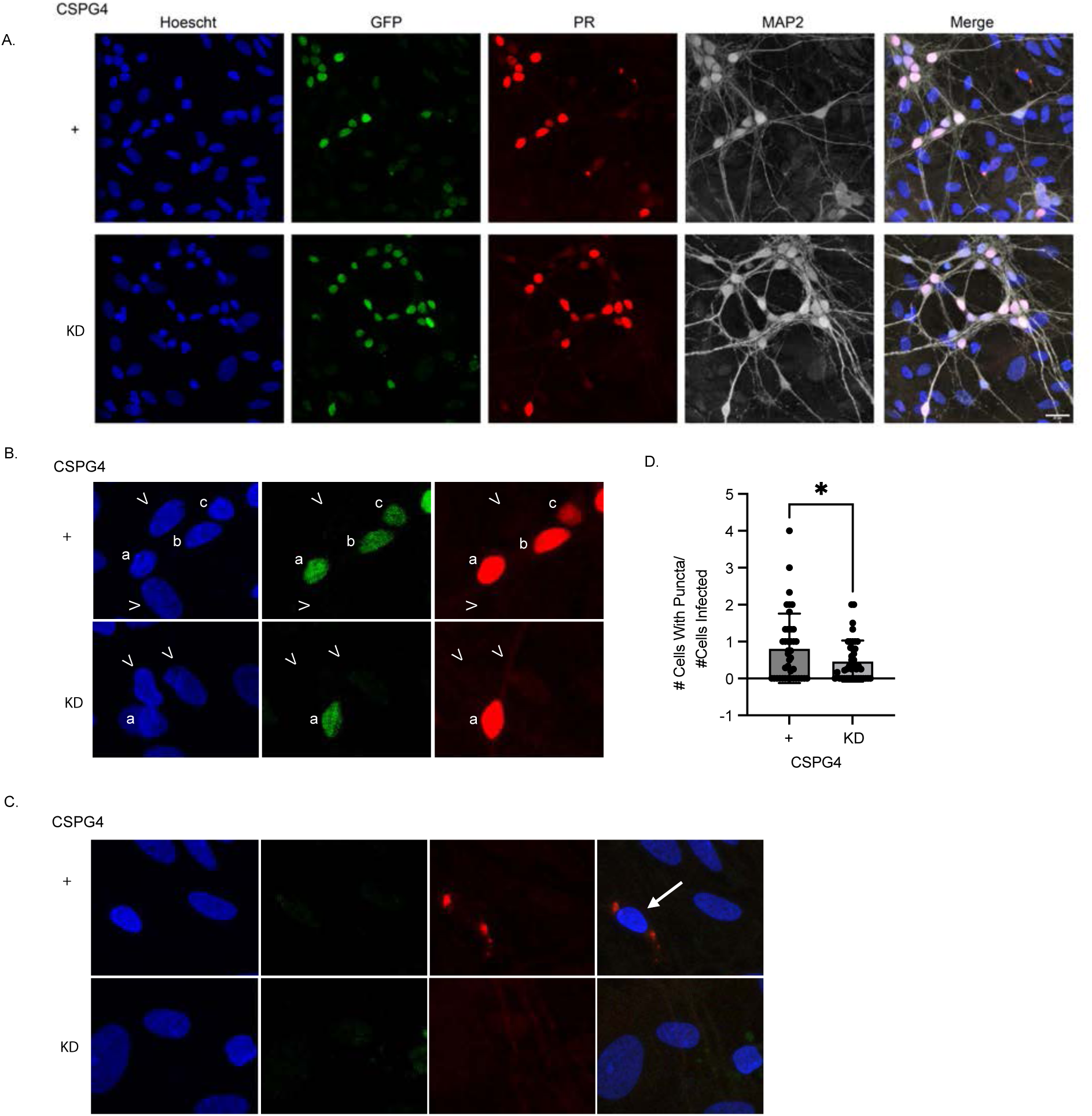
Intercellular transmission of poly-PR is reduced with CSPG4 siRNA knockdown. (A) Representative ICC of negative control and CSPG4 siRNA knockdown primary rat mixed spinal cord cultures transduced with AAV encoding GFP-T2A-PR_50_-FLAG for 5 days. PR-positive, GFP-negative puncta represent cell-to-cell transmission of poly-PR. Blue = Hoescht, Red = PR, Green = GFP, Grey = MAP2. Scale bar, 20µm. (B) Representative images of PR_50_– and GFP-positive cells denoted by letters. Carrots denote uninfected cells and are GFP– and PR_50_-negative. Red = PR, Green = GFP. (C) Representative images of PR_50_-positive, GFP-negative cells denoted by an arrow. Blue = Hoescht, Red = PR, Green = GFP. (D) Quantification of (A) was done by measuring the number of PR50-positive, GFP-negative recipient cells normalized to the number of PR50-positive, GFP-positive donor cells. Unpaired *t* test, two-tailed, n=4 biological replicates,*p<0.05.

We next assessed whether CSPG4 knockdown affected PR_50_ intercellular transmission. Viral transduction efficiency appeared comparable between CSPG4 knockdown and control siRNA-treated neurons (Fig. S5A). To quantify transmission, we measured the number of PR_50_-positive, GFP-negative recipient cells (Fig. 5C) normalized to the number of PR_50_-positive, GFP-positive donor neurons (Fig. 5B). CSPG4 knockdown significantly reduced the frequency of PR_50_ transmission, supporting its role in mediating neuron-to-neuron DPR propagation (Fig. 5D).

## Discussion

Here we investigated how arginine-rich DPRs, toxic products of the *C9orf72* hexanucleotide repeat expansion, transit between cells. We identified CSPG4 as a critical mediator of cellular uptake and intercellular transmission of arginine-rich DPRs. Although poly-PR and poly-GR have previously been shown to spread intercellularly (42), the mechanisms governing their cellular uptake and spread have remained unclear.

HA-PR_20_ was rapidly internalized through a temperature-dependent fluid-phase endocytic pathway that depends on CSPG4. CSPG4 is a GAG-modified cell surface protein, and under physiological conditions, carries a net negative charge. Since poly-PR and poly-GR are positively charged, our results supported a model in which electrostatic interactions initiate DPR internalization and subsequent cell-to-cell spread. Our findings suggested that after release from “donor” cells, poly-PR itself is exposed and then recognized by “recipient” cells. This was consistent with previous work that proposed an exosome-independent release of poly-PR (42).

Proteoglycans have been implicated in the uptake of other aggregation-prone proteins involved in neurodegeneration including Aβ (66), tau (67), and α-synuclein (67). Identification of CSPG4 through proximity labeling extends this concept to *C9orf72*-associated ALS/FTD, suggesting convergence of pathogenic mechanisms across multiple proteinopathies. Moreover, expression of CSPGs is frequently upregulated in response to nervous system injuries, including with neurodegeneration (68). One such example is the accumulation of CSPGs in the microenvironment surrounding spinal motor neurons in ALS transgenic rats (69).

While our findings indicated that CSPG4 is a key mediator for poly-PR uptake and intercellular spread, these conclusions were derived from cell lines and primary rat neuronal cultures. Future studies using *in vivo* models are necessary to determine whether this mechanism translates to the more complex brain microenvironment. Additionally, our experiments focused on DPRs containing 20 and 50 repeats, whereas patients may produce a broader spectrum of DPR species of various lengths. Given that proteoglycans frequently function as co-receptors (70, 71), it is also possible that additional cell surface proteins collaborate with CSPG4 to mediate DPR internalization. Elucidating these potential cooperations will be important for fully understanding arginine-rich DPR spread.

Insofar that DPR propagation contributes to disease progression, CSPG4 or its GAG chains could represent an intriguing therapeutic target. For example, small molecules or antibodies to inhibit or reduce CSPG4 levels in patients with *C9orf72*-mediated disease may be useful. Pharmacological inhibition of CSPG4-dependent uptake could block DPR internalization and intercellular transmission, thereby mitigating downstream effects such as nucleolar stress, nuclear transport defects, and impaired translation.

## Materials and Methods

### Antibodies

The following antibodies were used in this study: Clathrin Heavy Chain Monoclonal Antibody (X22) (ThermoFisher MA1-065), Dynamin 1 Monoclonal Antibody (ThermoFisher 3G4B6), PAK1 (Cell Signaling #2602), HA-Tag (6E2) Mouse mAb (Cell Signaling #2367), HA-Tag (C29F4) Rabbit mAb (Cell Signaling #3724), GAPDH (Sigma G8795), B-Actin (Novus NB600-501), Recombinant Anti-NG2 antibody [EPR23976-145] (Abcam ab275024), Anti-Fibrillarin antibody – Nucleolar Marker (Abcam ab5821), TDP-43 (Proteintech 10782-2-AP), Lamin A/C (4C11) Mouse mAb (Cell Signaling #4777), Rab7 (D95F2) XP® Rabbit mAb (Cell Signaling #9367), Purified anti-Neurofilament H (NF-H), Nonphosphorylated Antibody (Fisher 50-102-9517), poly(PR) (L. Petrucelli Rb8736), anti-MAP2 chicken polyclonal antibody (Abcam ab5392), GFP (B-2) (Santa Cruz Biotechnology NC9830411), FLAG (Cell Signaling 2368S), Invitrogen™ Wheat Germ Agglutinin, Alexa Fluor™ 594 Conjugate (Fisher W11262).

### Peptides

Peptides with the following sequences were generated with 98% purity through CASLO ApS: 1) (PR)_20_YPYDVPDYA; and 2) (GR)_20_YPYDVPDYA. “D” amino acid HA-PR_20_ was generated through GenScript. Lyophilized peptide was resuspended with sterile water to a stock concentration of 1 mM. Peptide was aliquoted and flash frozen with liquid nitrogen until use at final concentration of 2 μM.

### Silencer Select siRNAs

The following siRNAs were used in this study: Negative control #1 (human, rat; 4390843), CLTC (human s475), DNM2 (human s4212), PAK1 (human s10019), CSPG4 (human s3653), CSPG4 (rat s135455), PCDH7 (human s10109), BSG (human s2099), THBS1 (human s14099), CELSR3 (human s4524), APP (human s1500), CD44 (human s2683), LAMA4 (human s8062), PLAUR (human s10615), LAMB1 (human s535145), NOTCH2 (human s532526), EVC (human s4871), ACTN4 (human s959), CELSR2 (human s4527), SDC2 (human s12637), VCL (human s14764), PCDHAC2 (human s31972), ACTN1 (human s965), NRP1 (human s16844), PLPP2 (human s16382), XYLT2 (human s34495),

### Cell Lines and Cell Culture

All cells were maintained at 37°C in 5% CO_2_. HeLa cells were cultured in DMEM (Gibco™ 11965092) supplemented with 10% FBS (Fisher FB12999102) and 1% penicillin-streptomycin (Gibco™ 15140122). HEK293 Flp-In T-REx (ThermoFisher R78007) cells were maintained in DMEM supplemented with 10% Tetracycline free FBS (Gibco™ A4736201), 1% penicillin-streptomycin, 1% Glutamax (Gibco™ 35050061), 15μg/mL Blasticidin S (Gibco™ A1113903) and 100ug/mL zeocin (Gibco™ R25001). HEK293 Flp-In T-REx CSPG4-inducible stable cell lines were maintained in DMEM supplemented with 10% Tetracycline free FBS, 1% penicillin-streptomycin, 1% Glutamax, 15 μg/mL Blasticidin S, and 150 ug/mL Hygromycin B (Gibco™ 10687010). Primary rat mixed spinal cord cultures were prepared as previously described in (72).

### CSPG4^−/−^ CRISPR cell line creation

CSPG4^−/−^ cells were generated in the HEK293 Flp-In T-REx cell line. crRNA guide (IDT Hs.Cas9.CSPG4.AA and Hs.Cas9.CSPG4.AC) and tracrRNA (IDT 1073189) duplexes were combined with Cas9 nuclease (IDT 1081058) and transfected with RNAiMAX lipofectamine. Single colonies were isolated and expanded. Direct PCR lysis (Viagen 301-C) was performed to isolate genomic DNA which was subsequently sequenced to identify successful edits.

### CSPG4 Flp-In T-REx cell line generation

*plasmid generation:* pcDNA5-CSPG4 was cloned using standard methods. *stable cell line generation:* For CSPG4 stable cell line generation, 3×10^5^ cells of either HEK293 Flp-In T-REx or CSPG4^−/−^ HEK293 Flp-In T-REx were plated per well in a 6-well plate. Approximately 48 hours later, at 70% confluency, cells were transfected with 125 ng of CSPG4-pcDNA5/FRT/TO, 1.125 μg of pOG44 (Invitrogen™ V600520), and 6 μL of Lipofectamine 2000 (Invitrogen™ 11668500) per well. For selection purposes, cells were re-plated in media lacking zeocin but containing 150 μg/mL Hygromycin B. Cells were screened for successful integration via western blotting. *Doxycycline induction:* 6-well plates were coated for 30 minutes with rat tail collagen. Plates were rinsed twice with PBS, and 1.2×10^6^ doxycycline-inducible cells were plated per well. 24 hours later, CSPG4 expression was induced with 1.25 pg of doxycycline for 24 hours. Cells were either collected for CSPG4 expression or pulse/chased with DPR.

### Inhibitors

*Endocytosis Inhibitors:* Cells were pretreated with endocytosis inhibitors for 1 hour (Chlorpromazine: 10 μg/mL, EIPA: 100 μM, Dynole 31-2 and 34-2: 5 μM). Then 2 μM DPR was applied either through pulse/chase or bath application. *CSPG Inhibitors:* Cells were pretreated with Chondroitinase ABC from Proteus vulgaris (Millipore-Sigma C2905-5UN) (100 μM HeLa/HEK and 20-50 μM Neurons) for 1 hour before pulse/chase or bath application with DPR. Primary rat mixed spinal cord cultures were pretreated with 10 μM Fluorosamine or Difluorosamine for 24 hours before a 2-hour bath application of 2 μM HA-PR_20_.

### siRNA KD

*HeLa:* Cells were reverse transfected with silencer select siRNAs and RNAiMAX lipofectamine. 3×10^5^ cells were plated per well of a 6-well plate. 48 hours later, downstream assays were performed. *Primary rat neurons:* primary rat cortical neuronal cultures and primary rat mixed spinal cord cultures were transfected on DIV4 with silencer select siRNAs and RNAiMAX lipofectamine. Cultures were maintained until DIV14 when downstream assays were performed.

### Pulse/Chase Assay

Protocol was adapted from (73). 6×10^5^ HeLa cells per well were plated in a 6-well plate. 24 hours later, cells were pulsed with 2 μM HA-PR_20_ or HA-GR_20_ for 5 minutes at 37 °C. Then, cells were washed with PBS and replaced with fresh media at 37 °C for 15-minutes. Cells were washed 3 times with PBS and incubated with trypsin for 5-minutes at 37°C. Enzyme inhibitors (100 μL (one Complete Mini tablet, Roche, in 2.5mL PBS) and BSA (100 μL of 1 mg/mL) were added. Then samples were spun down at 500xg for 5 minutes. Samples were washed with wash buffer (1 mL 50 mM Tris buffer pH 7.4, 0.1% BSA) and spun down at 500xg for 5 minutes. Next, samples were resuspended in lysis buffer (50 mM Tris pH 7.4, 1 M NaCl, 1% NP40) and sonicated at 20% for 10 seconds each before spinning down at max speed for 10 minutes. Supernatant was collected and 20 μg of protein was loaded per well for western blotting analysis.

### Bath Application

2 μM HA-PR_20_ or HA-GR_20_ was added to media for various amounts of time. Then cells were either fixed for immunocytochemistry analysis or lysed for immunoblotting analysis.

### Temperature Shift Assay

1×10^5^ HeLa cells were plated per well on poly-lysine coated coverslips in a 12-well plate. 24 hours later, the cells were acclimated on ice or at room temperature for 30 minutes. While on ice, a 5-minute pulse with 2 µM PR_20_-HA was performed. Cells were washed once with PBS and twice with 0.1 M glycine, pH 3.0 for five minutes to remove unbound peptide. Fresh media was added, and cells were shifted to a 37 °C incubator for 30 minutes before fixing.

### Immunocytochemistry

Cells grown on poly-lysine coated coverslips (Azer Scientific 200181) were fixed with 4% paraformaldehyde for 15 minutes at RT and washed three times with PBS. Then, cells were permeabilized with 0.1% Triton-X 100 (Sigma T8787) in PBS for 10 minutes at RT and washed three times with PBS. Samples were blocked for 30 minutes at RT with 2% BSA (Sigma 10735086001) in PBS. Coverslips were incubated in primary antibody diluted in blocking solution overnight at RT. The next day, samples were washed twice with PBS and incubated in Alexa Fluor secondary antibody (1:500) and Hoescht (1:1000) diluted in blocking solution for 1-4 hours at RT. Coverslips were washed twice with PBS and once with milliQ water before mounting on slides (Fisher 23888100) using Electron Microscopy Sciences Fluoro-Gel Mounting Medium with Tris Buffer (Fisher 50-247-04). Coverslips were sealed onto slides with nail polish before imaging.

### Confocal Microscopy

Confocal microscopy was performed using a Leica DMI4000B Microscope. Once laser intensities and photomultiplier levels were set, data was collected with no further adjustments for all data collected per experiment. Neuronal ICC experiments were imaged and quantified in a blinded manner. Relative Nuclear HA signal was achieved by normalizing HA nuclear corrected total cell fluorescence (CTCF) to Hoescht nuclear CTCF in ImageJ.

### Structures of Reagents Used for µMap Experiments

Diazirine-biotin and Ir-catalyst DBCO were prepared as described previously(74, 75).

### Preparation of PR_20_-Ir-HA Peptide

Preparation of PR_20_-Ir-HA was performed with an adapted protocol from Seath et al(74). (PR)_20_-(MMT-protected C)-HA was synthesized on a 0.1 mmol scale by standard Fmoc solid-phase peptide synthesis using DIC-Oxyma activation on a CEM Liberty Blue microwave-assisted peptide synthesizer on ChemMatrix Rink amide resin. Each residue was double coupled during the synthesis. Fmoc deprotection was performed at room temperature by the addition of 20% piperidine in dimethylformamide (DMF) with 0.1 M hydroxybenzotriazole. Treatment was performed twice at room temperature for 30 min. The peptide resin was stored at 4°C before proceeding to the iridium catalyst conjugation step.

For preparation of Maleimide-PEG3-DBCO/N3-Ir, 124.1 µL of Azido-PEG3-NHS ester (Broadpharm) (50 mg/mL freshly prepared stock in DMF), 91.6 µL of 2-aleimidoethylamine trifluoroacetate salt (Broadpharm) (50 mg/mL stock in DMF), and 102.5 µL of DIPEA solution (50 mg/mL stock in DMF) were combined and incubated at room temperature for 1 h. 29.1 mg Ir-catalyst DBCO (2) was added to the solution. The solution was vortexed and incubated for 1 h at room temperature.

In a separate 1.5 mL Eppendorf tube, for MMT deprotection, 50 mg of resin was deprotected with 10 x 0.5 mL x 2 min washes of 1% v/v TFA and 5% v/v TIPS in DCM. The resin was washed 2 x with 500 µL with DMF, followed by 3 x 500 µL 0.3% v/v DIPEA in DMF. The pH of the final slurry was checked via pH paper to ensure basicity.

24.3 µL of DIPEA solution (50 mg/mL stock in DMF) was added to the solution containing Maleimide-PEG3-DBCO/N3-Ir, followed by 400 µL of DMF. The resulting solution was added to the resin, and the solution was bubbled using a steady stream of nitrogen and a needle for 2 h. The resin was washed with 3 x 1 mL of DMF, followed by 3 x 1 mL of DCM. The resin was dried using a vacuum manifold and transferred to a 15 mL conical tube. Cleavage from the resin and side chain deprotection was performed by adding 1000 µL of 92.5:5:2.5 TFA:thioanisole:ethanedithiol. The cleavage reaction was incubated on a mechanical shaker at 250 rpm. After 3 h, the solution was filtered, and solvent was removed using a steady stream of nitrogen. The cleaved peptide was precipitated using cold diethyl ether and isolated by refrigerated centrifugation. The crude peptide was dissolved in 1 mL DMSO, purified by preparative scale reversed-phase high-performance liquid chromatography, and characterized by mass spectrometry.

### Target Identification of PR_20_ via µMap Proximity Labeling

*Preparation of PR*_20_*-Ir-HA solution:* From a 100 µM PR_20_-Ir-HA concentrated stock in DMSO, PR_20_-Ir-HA was diluted to a final concentration of 200 nM in DPBS supplemented with an additional 100 mM NaCl. The solution was warmed in a 37 °C water bath for 1 h protected from light.

*PR*_20_ *µMap Proximity Labeling at 5 min timepoint:* 12 x 10 cm plates of HeLa cells were grown to near confluency in DMEM media supplemented with 10% FBS and 100 U/mL penicillin-streptomycin at 37°C with 5% CO_2_. The plates were removed from the incubator and acclimated on ice for 30 minutes under ambient atmosphere. The media was removed, and the cells were washed twice with 2 mL of 4 °C DPBS. 3 mL of the PR_20_-Ir-HA solution was added to cells, and the cells were incubated on ice for 5 minutes. The cells were washed once with 2 mL of 4 °C DPBS, and 3 mL of 250 µM diazirine-biotin (**1**) in 4 °C DPBS was added. The plate was irradiated with 450 nm light for 3 minutes at 100% intensity using an Efficiency Aggregators Bio-Photoreactor BPR200. The diazirine solution was aspirated, and the cells were washed twice with 2 mL of 4 °C DPBS before proceeding to the *Dissociation of cells from plates and membrane fractionation* protocol.

*PR*_20_ *µMap Proximity Labeling at 1 hour timepoint*: 12 x 10 cm plates of HeLa cells were grown to near confluency in DMEM media supplemented with 10% FBS and 100 U/mL penicillin-streptomycin at 37 °C with 5% CO_2_. The plates were removed from the incubator and acclimated on ice for 30 minutes under ambient atmosphere. The media was removed, and the cells were washed twice with 2 mL of 4 °C DPBS. 3 mL of the PR_20_-Ir-HA solution was added to cells, and the cells were incubated on ice for 5 minutes. The cells were washed once with 2 mL of 4 °C DPBS, and 3 mL of 37 °C complete DMEM was added. Cells were incubated at 37 °C with 5% CO_2_ for 1 h. The media was aspirated, and 3 mL of 250 µM diazirine-biotin (1) in 4 °C DPBS was added. The plate was irradiated with 450 nm light for 3 minutes at 100% intensity using an Efficiency Aggregators Bio-Photoreactor BPR200. The diazirine solution was aspirated, and the cells were washed twice with 2 mL of 4 °C DPBS before proceeding to the *Dissociation of cells from plates and membrane fractionation* protocol.

*Dissociation of cells from plates and membrane fractionation:* After aspirating, 1 mL of 4 °C DPBS was added to the plate. Cells were dissociated using a cell scraper and transferred to a 15-mL conical tube. The plate was rinsed with an additional 1 mL of 4 °C DPBS, and the solution were combined into the same 15 mL conical tube. Cells from 4 plates were combined into a single 15 mL conical tube for each replicate (n=6).

The cells were centrifuged at 1,000xg for 5 min at 4 °C. The supernatant was removed, and the cell pellets were frozen at –20 °C overnight. The next day, the samples were thawed on ice, and each sample was resuspended in 1 mL of membrane permeabilization buffer (MEM-PER Plus Membrane Fractionation Kit, Thermo Fisher Scientific, 89842) containing 1X protease inhibitors (Roche, 11697498001) and incubated for 20 min at 4 °C. The samples were centrifuged at 16,000xg for 15 min at 4°C. The supernatant enriched in cytosolic proteins was removed, and the pellet was resuspended in 300 μL RIPA containing 1% SDS and 1X protease inhibitors. The samples were placed on ice and sonicated using a Branson tip sonicator to break up the membrane pellet once for 5 seconds at 35% intensity and heated for 5 min at 95°C. The samples were diluted to 1.0 mL with RIPA, transferred to 2.0 mL tubes, and sonicated twice for 5 seconds at 35% intensity with a 30 seconds off cycle. The protein concentration was measured by BCA assay before proceeding to the *Streptavidin Bead Enrichment and Proteomics Sample Preparation* protocol.

### Streptavidin Bead Enrichment and Proteomics Sample Preparation

For streptavidin bead enrichment, 800 µg of membrane cell lysate was added to a 1.5 mL LoBind tube containing 100 μL of Pierce streptavidin magnetic beads (Thermo Fisher Scientific, 88817) that were pre-washed twice with 1 mL RIPA buffer. The samples were incubated overnight on a rotator at 4 °C, and the beads were pelleted on a magnetic rack. The supernatant was removed, and the beads were washed 3x with 1 mL 1% SDS in DPBS, 3x with 1 mL 1M NaCl in DPBS, and 3x with 1 mL 10% EtOH in DPBS. The beads were incubated in each of the washes for 5 min on a rotator before pelleting to remove the wash. The beads were pelleted to remove supernatant, resuspended in 0.5 mL RIPA, and transferred to a new 1.5 mL LoBind tube.

The supernatant was removed, and the beads were washed with 3 x DPBS (0.5 mL) and 3 x NH4HCO3 (0.5 mL) (100 mM in water), incubating 5 min on a rotator for each wash. The beads were resuspended in 500 μL 3 M urea in DPBS and 25 μL of 200 mM DTT in 25 mM NH4HCO3 was added. The beads were incubated at 55 °C for 30 min on a rotator. Subsequently, 30 μL 500 mM iodoacetamide in 25 mM NH4HCO3 was added and incubated for 30 min on a rotator at room temperature in the dark. The supernatant was removed, and the beads were washed with 3 x 0.5 mL DPBS and 3 x 0.5 mL triethyl ammonium bicarbonate (TEAB, 50 mM), incubating 5 min on a rotator for each wash. The beads were resuspended in 0.5 mL TEAB (50 mM) and transferred to a new protein LoBind tube. The beads were resuspended in 40 μL TEAB (50 mM), 2.0 μL trypsin (1 mg/mL in 50 mM acetic acid) was added, and the beads were incubated overnight with end-over-end rotation at 37 °C. After 16 hours, an additional 1.0 μL trypsin was added and the beads were incubated for an additional 1 hour at 37 °C. Meanwhile, TMT 6-plex label reagents (0.8 mg) (Thermo) were equilibrated to room temperature and diluted with 41 μL of anhydrous acetonitrile (Optima grade) and centrifuged. The beads were pelleted, and the supernatant was added to the appropriate TMT 6-plex tube. The reaction was incubated for 2 hours at room temperature. The samples were quenched with 8 μL of 5% hydroxylamine in water and incubated for 15 minutes. The samples were pooled in a new Protein LoBind tube and quenched with TFA (16 μL, Optima). TMT-labeled peptides were dried down in a SpeedVac, re-dissolved in 300 μL of 0.1% TFA in water and fractionated into 8 fractions using the PierceTM High pH Reversed-Phase Peptide Fractionation Kit (#84868). Fractions 1, 4, and 7 were combined as sample 1. Fractions 2 and 6 were combined as sample 2. Fractions 3, 5, and 8 were combined as sample 3. Three combined samples were dried completely in a SpeedVac and resuspended in 20 μL 5% acetonitrile/water (0.1% formic acid (pH = 3)). 2 μL (∼ 360 ng protein) was injected per run using an Easy-nLC 1200 UPLC system. Samples were loaded directly onto a 45cm long x 75 μm inner diameter nano capillary column packed with 1.9 μm C18-AQ resin (Dr. Maisch, Germany) mated to a metal emitter in-line with an Orbitrap Fusion Lumos (Thermo Scientific, USA). The column temperature was set at 45°C, and a two-hour gradient method running at a flow rate of 300 nl min-1 was used. The mass spectrometer was operated in data dependent mode with synchronous precursor selection (SPS) – MS3 method with 120,000 resolution MS1 scan (positive mode, profile data type, intensity threshold 5.0e3 and mass range of 375-1600 m/z) in the Orbitrap followed by CID fragmentation in the ion trap with 35% collision energy for MS2 and HCD fragmentation in the Orbitrap (50,000 resolution) with 55% collision energy for MS3. The MS3 scan range was set to 100-500 with injection time of 120 ms. A dynamic exclusion list was invoked to exclude previously sequenced peptides for 60 s and a maximum cycle time of 2.5 seconds was used. Peptides were isolated for fragmentation using the quadrupole (0.7 m/z isolation window). The ion-trap was operated in Rapid mode.

### Proteomics Data Processing and Quantitation for TMT Data (FragPipe)

Files were converted from.raw files to mzmL using the following protocol (https://fragpipe.nesvilab.org/docs/tutorial_convert.html). Quantitation was performed using the TMT10-MS3 workflow in FragPipe v19 (MSFragger 3.7, IonQuant-1.8.10, philosopher 4.8.0, python 3.9.12)50 with all eight fractions set as a single experiment. The Homo sapiens (Human) fasta from Uniprot (UP000005640) with added decoys and common contaminants was used as a database. The resulting matrix.pg file was opened in Perseus (v2.0.7.0). Intensities were inputted as “main”, the rest of the descriptors are categorical. Data was then transformed (Log base 2). Data was annotated by sample groups. Normalization was performed via median subtraction. Following this process, a volcano plot was generated utilizing a t-test for statistical significance. Resulting volcano plots were plotted in GraphPad Prism 9 for visualization.

### Proximity Ligation Assay (PLA)

2 μM HA-PR_20_ was bath-applied to cells plated on coverslips for 30 minutes the day after plating. Cells were fixed with 4% Paraformaldehyde for 15 minutes. PLA was performed using Duolink® PLA Fluorescence Protocol (Sigma-Aldrich). Negative controls consisted of HA and CSPG4 antibodies alone.

### Sarkospin Fractionation

Sarkospin fractionation was carried out as described in (64, 76). 2 wells of a 6-well plate of HEK293 or rat primary cortical neuronal cultures were collected in PBS, pelleted by centrifugation, and resuspended in lysis buffer containing 0.5% sarkosyl and benzonase. Protein concentration was equalized before proceeding with sarkosyl solubilization and centrifugation to isolate sarkosyl insoluble material.

### Nuclear/Cytoplasmic Fractionation

2 μM HA-PR_20_ was bath applied per well to DIV17 primary rat cortical neurons growing in six well plates for 48 hours. Three wells were pooled and spun down in PBS. Then fractionation was performed using the NE-PER Nuclear and Cytoplasmic Extraction Kit (ThermoFisher 78833). Fractionation was assessed via immunoblotting, and 30 μg of protein was loaded per lane. Quantification was done by dividing Nuclear TDP-43 by Cytoplasmic TDP-43.

### Motor Neuron Survival Assay

On DIV14, mixed spinal cord cultures were treated with 2 μM HA-PR_20_ for 5 days before fixing. Coverslips were stained with SMI-32 antibody, and motor neurons were counted. Data collected was blind to experimental manipulation.

### AAV Transduction

Primary rat mixed spinal cord cultures growing on coverslips in 12 well plates were transduced with 5 μL/well of AAV encoding GFP-T2A-PR_50_-FLAG on DIV14. The next day, a full media change was done. Five days after transduction, coverslips were fixed and immunocytochemistry was performed. AAV encoding GFP-T2A-PR_50_-FLAG was prepared by Vector Builder.

### Statistical Analysis

Microsoft Office Excel and GraphPad Prism were used for data analysis and to plot graphs. Data were expressed as mean ± standard deviation. For comparisons among two groups, two-tailed unpaired Student’s t-tests were used, unless specifically noted. For comparisons of more than two groups, ordinary one-way ANOVA with Tukey’s *post hoc* test was used. P ≤ 0.05 was considered significant. All data presented is derived from no less than 3 experimental replicates. For experiments with no technical replicates, the control is expressed as 1 and the experimental group is expressed relative to control. If the experiment had technical replicates, each data point within the biological replicate is expressed relative to the average of the control technical replicates.

## Acknowledgements

A.B.S., J.M.P., C.D. and R.G.K. are supported by the US Public Health Service (NS122908, NS124802), the US Department of Defense (W81XWH-21-1-0236), the Les Turner ALS Foundation, the Kissick Family Foundation and the Milken Institute Science Philanthropy Accelerator for Research and Collaboration and the Heather Koster Family Charitable Fund. B.F.B., N.A.T., D.C.M., and D.W.C.M. are supported by the National Institute of General Medical Sciences of the National Institutes of Health (R35GM134897) and the Princeton Catalysis Initiative. We thank members of the Kalb laboratory for technical assistance, critical discussions and project guidance during the undertaking of this work. We thank Dr. Petrucelli for kindly providing the poly(PR) antibody. We are grateful to Dr. Wong and Dr. Ling for the generous gift of Fluorosamine and Difluorosamine CSPG synthesis inhibitors.

## Author Contributions

A.B.S, B.F.B, J.M.P., C.D., N.A.T., D.W.C.M, R.G.K. designed research. A.B.S., B.F.B., J.M.P., C.D., N.A.T., D.C.M., performed research. A.B.S., B.F.B., J.M.P., C.D., D.C.M. analyzed data. A.B.S. and R.G.K. wrote the paper.

## Competing Interest Statement

The authors have no competing interests to disclose.

## Notes

### Competing Interest Statement

The authors have declared no competing interest.

