## Supplemental Figures for "The molecular mechanism of uptake and cell-to-cell transmission of arginine-containing dipeptide repeat proteins"

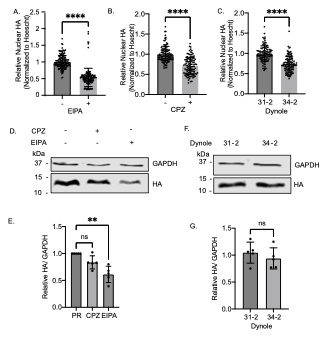

**Fig. S1.** Uptake of PR_20_ in HeLa cells is mediated by a pathway that shares features with macropinocytosis. (A) Quantification of nuclear HA in HeLa cells after pretreatment with 5-(N-ethyl-N-isopropyl) amiloride (EIPA), an inhibitor of macropinocytosis, and 30-minute bath application of HA-PR_20_. Unpaired *t* test, two-tailed, n=3 biological replicates, ****p<0.0001. (B) Quantification of nuclear HA in Hela cells after pretreatment with chlorpromazine (CPZ), an inhibitor of clathrin-mediated endocytosis, and 30-minute bath application of HA-PR_20_. Unpaired *t* test, two-tailed, n=3 biological replicates, ****p<0.0001. (C) Quantification of nuclear HA in Hela cells after pretreatment with dynamin inhibitor Dynole 34-2 or its inactive isomer Dynole 31-2, and 30-minute bath application of HA-PR_20_. Unpaired *t* test, two-tailed, n=3 biological replicates, ****p<0.0001. (D) Representative western blot of internalized HA from HeLa cells pretreated with CPZ or EIPA followed by a 5-minute pulse with HA-PR_20_ and a 5-minute chase. GAPDH, loading control. (E) Quantification of (D). One way ANOVA with Tukey’s *post hoc* test, F _(2,12)_ = 15.11, n=5 biological replicates, **p<0.01. (F) Representative western blot of internalized HA from HeLa cells pretreated with Dynole 31-2 or 34-2 followed by a 5-minute pulse with HA-PR_20_ and a 5-minute chase. GAPDH, loading control. (G) Quantification of (F). Unpaired *t* test, two-tailed, n=5 biological replicates, ns.

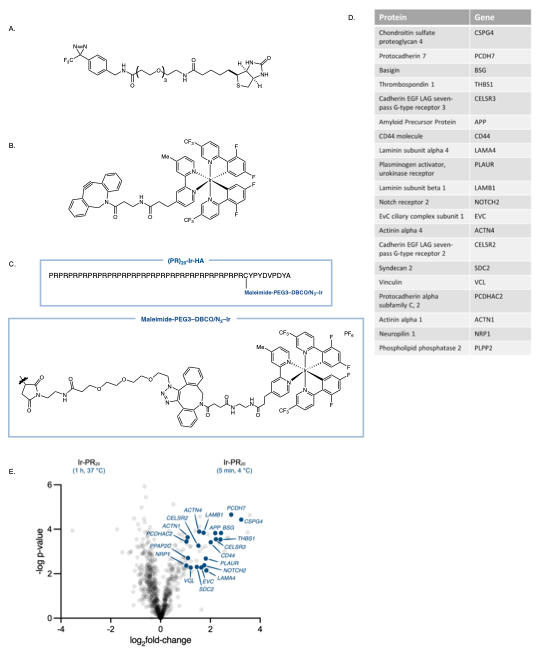

**Fig. S2.** Candidate mediators of PR_20_ uptake were obtained through µMap proximity labeling technique. (A) Diazirine-biotin. (B) Ir-catalyst DBCO. (C) Preparation of PR_20_-Ir-HA Peptide. (D) Filtered list of twenty candidates resulting from µMap followed by mass spectrometry analysis. Candidates were filtered based on two criteria: (1) expression in neurons, and (2) localization to the cell surface. These twenty candidates were selected for an siRNA-based screen in HeLa cells using our pulse-chase assay to quantify internalized HA-PR_20_ by western blotting. (E) Volcano plot of top 20 protein candidates.

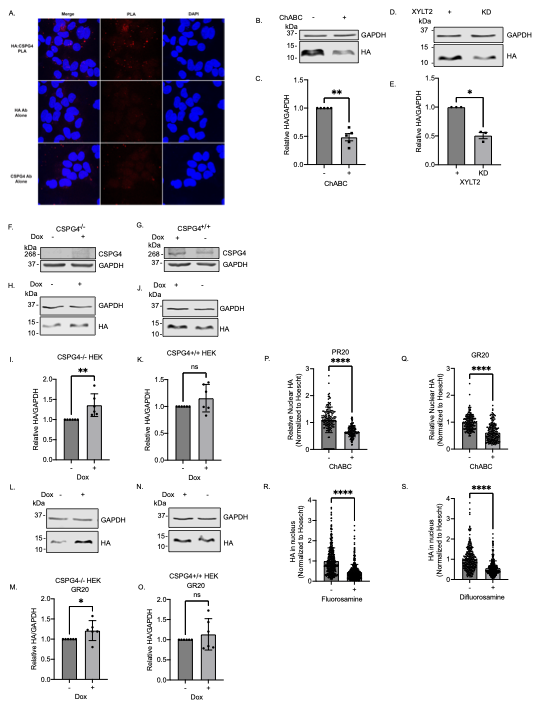

**Fig. S3.** Inhibiting CSPGs reduces uptake of arginine-rich DPRs. (A) Representative ICC of HA-PR_20_:CSPG4 Proximity Ligation Assay (PLA) in HEK cells after 30-minute HA-PR_20_ bath application. Red signal indicates proteins are within 40nm. Each antibody alone was used as negative controls. Blue = Hoescht. (B) Representative western blot of a 5-minute HA-PR_20_ pulse followed by a 5-minute chase in HeLa cells after pretreatment with Chondroitinase ABC (ChABC), an enzyme that digests chondroitin sulfate glycosaminoglycan (GAG) chains. GAPDH, loading control. (C) Quantification of (B). Unpaired *t* test, two-tailed, n=5 biological replicates, **p<0.01. (D) Representative western blot of a 5-minute HA-PR_20_ pulse followed by a 5-minute chase in HeLa cells after Xylosyltransferase 2 (XYLT2) siRNA knockdown. XYLT2 is an enzyme responsible for initiating GAG chain addition to proteoglycans. GAPDH, loading control. (E) Quantification of (D). Unpaired *t* test, two-tailed, n=3 biological replicates, *p<0.05. (F) Representative western blot of CSPG4 dox induction in CSPG4 dox-inducible CSPG4^-/-^ Flp-In T-REx HEK cells. GAPDH, loading control. (G) Representative western blot of CSPG4 dox induction in CSPG4 dox-inducible Flp-In T-REx HEK cells. GAPDH, loading control. (H) Representative western blot of a 5-minute HA-PR_20_ pulse followed by a 15-minute chase in CSPG4 dox-inducible CSPG4^-/-^ Flp-In T-REx HEK cells -/+ dox. GAPDH, loading control. (I) Quantification of (H). Unpaired *t* test, two-tailed, n=6 biological replicates, **p<0.01. (J) Representative western blot of a 5-minute HA-PR_20_ pulse followed by a 15-minute chase in CSPG4 dox-inducible Flp-In T-REx HEK cells -/+ dox. GAPDH, loading control. (K) Quantification of (J). Unpaired *t* test, two-tailed, n=6 biological replicates, ns. (L) Representative western blot of a 5-minute HA-GR_20_ pulse followed by a 15-minute chase in CSPG4 dox-inducible CSPG4^-/-^ Flp-In T-REx HEK cells -/+ dox. GAPDH, loading control. (M) Quantification of (L). Unpaired *t* test, two-tailed, n=6 biological replicates, **p<0.05. (N) Representative western blot of a 5-minute HA-GR_20_ pulse followed by a 15-minute chase in CSPG4 dox-inducible Flp-In T-REx HEK cells -/+ dox. GAPDH, loading control. (O) Quantification of (N). Unpaired *t* test, two-tailed, n=6 biological replicates, ns. (P) Quantification of nuclear HA-PR_20_ in neurons pretreated with Chondroitinase ABC before bath application of HA-PR_20_. Unpaired *t* test, two-tailed, n=3 biological replicates, p<0.001. (Q) Nuclear HA-GR_20_ quantification in neurons pretreated with Chondroitinase ABC before bath application of HA-GR_20_. Unpaired *t* test, two-tailed, n=3 biological replicates, ****p<0.001. (R) Nuclear HA-PR_20_ quantification in neurons pretreated with Fluorosamine, a CSPG inhibitor, before bath application of HA-PR_20_. Unpaired *t* test, two-tailed, n=3 biological replicates, ****p<0.001. (S) Nuclear HA-PR_20_ quantification in neurons pretreated with Difluorosamine, a CSPG inhibitor, before bath application of HA-PR_20_. Unpaired *t* test, two-tailed, n=3 biological replicates, ****p<0.001.

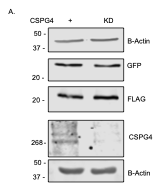

**Fig. S5.** Control and CSPG4 knockdown neurons transduced with GFP-T2A-PR_50_-FLAG both express GFP and PR_50_-FLAG. (A) Representative western blot of rat pure cortical neurons transduced with AAV encoding GFP-T2A-PR_50_-FLAG for 5 days in negative control and CSPG4 siRNA knockdown conditions. GFP and FLAG immunoblotting indicates that transduction levels are similar between both negative control and CSPG4 knockdown conditions. CSPG4 immunoblotting demonstrates CSPG4 knockdown level. B-Actin was used as a loading control.
